# Physiological limits of localized hypothermia in the human cochlea: The role of vascular heat transport

**DOI:** 10.64898/2026.07.14.738525

**Authors:** Bailey McCorkendale, Ricardo Rodriguez, Rowan Fink, MacKenzie Moore, Steven A. Romero, Fateme Esmailie

**Affiliations:** Department of Biomedical Engineering, University of North Texas, Denton, Texas, USA; Department of Physiology and Anatomy, UNT Health Fort Worth, Texas, USA

**Keywords:** Mild therapeutic hypothermia, Computational modeling, Blood perfusion, Bioheat transfer

## Abstract

**Purpose:** Mild therapeutic hypothermia (MTH) preserves cochlear function in animal models and is now entering early-phase human trials for hearing preservation. However, the extent to which the human cochlea can actually be cooled, and the mechanisms underlying MTH, remain unclear, in part because blood perfusion is expected to oppose localized cooling. In this study we evaluated the impact of blood flow on human cochlear temperature exposed to the MTH device using a combined experimental and computational approach.

**Methods:** Temperature measurements were obtained from a human cadaver skull exposed to a commercial MTH device. These data were used to validate a three-dimensional bioheat transfer model incorporating realistic skull anatomy. The validated model was subsequently extended to include physiological blood perfusion in the internal carotid artery; a major heat source located near the cochlea. Finally, the in silico model was further expanded to incorporate the surrounding skin and brain tissues.

**Results:** Incorporating blood flow in internal carotid artery substantially altered predicted cochlear temperature distributions, highlighting the importance of localized vascular heat transport in the human cochlea during MTH. Although cochlear cooling was attenuated in the presence of perfusion, the therapeutic effects of MTH may not depend solely on the magnitude of local intracochlear temperature reduction. Additional mechanisms, such as reduced facial surface temperature, may also contribute to its efficacy.

**Conclusion:** The validated in silico model provides a physiologically realistic framework for evaluating human cochlear thermal responses, investigating MTH mechanisms, and optimizing temperature-based strategies for hearing preservation.

## 1 Introduction

Hearing loss is one of the most common human medical conditions, arising from aging, congenital and genetic factors, and exposure to noise or ototoxic drugs. Hearing loss adversely affects life quality and may lead to depression, isolation, and cognitive decline [1, 2]. Currently, hearing aids and cochlear implants are common treatments for hearing loss [3, 4]. However, despite the success of hearing aids and cochlear implants in improving patients’ quality of life, these treatments face some challenges, including high cost [5], risk of damaging remaining hearing ability [6, 7], limitation in restoring all features of natural sound perception [8], and risk of post-operation infection [9]. Hearing loss prevention mitigates these challenges and can be used to provide individuals with a sustainable quality of life.

Noise-canceling or hearing protection devices (HPDs) [10, 11, 12, 13], medication [14], controlled temperature change (i.e., hypothermia and hyperthermia) [6, 7, 15, 16], and photobiomodulation (PBM) [17, 18, 19, 20], have been studied as possible methods to prevent hearing loss. Hearing protection devices (HPDs) are frequently disregarded in practice in part because they can impair situational awareness, thus making them less acceptable as routine personal protective equipment and contributing to poor user compliance [10, 11, 12, 13]. Hearing prevention medications are antioxidant-based therapies that eliminate free radical accumulation, but normal free radical levels are essential for biological functions and cellular hemostasis. Therefore, the use of drugs to eliminate hearing damage is an invasive method that can lead to interruption in biological functions [6, 21, 22, 23]. Temperature increase (Hyperthermia) [24] or temperature decrease (Hypothermia) [6, 7, 15, 16, 21, 25] are thermal methods applied to preserve hearing when there is a risk of hearing damage due to exposure to ototoxicity, noise, or cochlear implants excessive insertion force. Photobiomodulation, which involves the application of near-infrared light, has also emerged as a promising neuroprotective strategy for hearing preservation, although light-induced thermal effects must be carefully considered. Hyperthermia, hypothermia and PBM are non-invasive approaches with potential for clinical translation to support the maintenance of auditory function.

Among the three non-invasive methods for hearing preservation, hypothermia has shown the greatest success in neurotrauma treatment applications, particularly in preventing hearing loss [26, 27, 28]. Hypothermia is also the only thermal non-invasive hearing preservation strategy to have advanced into human clinical trials, using the RestorEar devices (ReBound Lite, ReBound, and ReBound Aero; ClinicalTrials.gov NCT06375278 and NCT06729632); however, its efficacy for hearing preservation in humans has not yet been established. The potential mechanism of action of hypothermia to mitigate hearing loss includes reduced uptake of ototoxic drugs, reduced inflammation, reduced reactive oxygen species (ROS) / reactive nitrogen species (RNS) formation, reduced apoptosis, and preservation of ribbon synapses on the inner hair cells [23].

Current studies on mild therapeutic hypothermia have primarily focused on in-vivo animal models, such as rats [26, 27, 29], ex-vivo studies on cadaveric temporal bone [6, 28] and cadaveric human head [13]. However, effective translation of findings from animal models to humans requires careful consideration of interspecies differences, including tissue thermophysical properties, blood perfusion dynamics, and anatomical factors such as cochlear and skull size. In the MTH study conducted by Rajguru et al., on rats, an active cooling device was applied using a fluid flow at a constant temperature of 5 *^◦^*C and achieved a maximum cochlear tissue temperature reduction of 6 *^◦^*C over 120 minutes [30]. This cooling protocol effectively led to preventing hearing loss without causing tissue damage [30]. The protocol involved a 12-minute induction (fluorocarbon temperature decreasing from 33*^◦^*C to 5*^◦^*C), followed by 2 hours of sustained cooling, and concluding with a 12-minute rewarming phase [30]. However, this cooling protocol is not achievable with the current MTH device (RestorEar) for human use (See section 3. Results).

A significant temperature gradient between the location of the RestorEar (MTH device) on the head and the middle and inner ear was reported in the recent cadaveric study by Yepes et al. [13]. They showed that RestorEar devices reduced inner ear temperature by 2.9*^◦^*C (Δ*T* = *−*2.9*^◦^*C) after 30 minutes, notably less than the 6*^◦^*C achieved in animal studies [13]. While these measurements provide valuable insight into human cochlear cooling performance, cadaver studies do not capture physiological processes such as blood perfusion and thermoregulatory responses, which are important determinants of thermal transport in vivo. In addition, direct measurement of cochlear temperature in living humans is not feasible due to ethical and safety considerations. Furthermore, the proximity of major blood vessels, including the carotid arteries and jugular veins, to both the cooling region and the cochlea may substantially influence heat transfer during hypothermia. Therefore, computational modeling provides a practical approach for investigating the role of blood perfusion and vascular heat transport on intracochlear temperature during MTH.

In this study, we evaluated the impact of blood perfusion on temperature reduction in human cochlea exposed to mild therapeutic hypothermia. We first experimentally examined the impact of RestorEar devices on cochlear temperature within a cadaver skull in the absence of blood perfusion and validated a computer model using the experimental data. We then modeled the effect of blood perfusion on cochlear temperature in full human head using an in silico model.

## 2 Materials and Methods

To evaluate the thermal response of the human cochlea to mild therapeutic hypothermia, we developed a multiscale computational framework and validated it using experimental measurements. We first reconstructed the human skull geometry and simulated temperature changes induced by an MTH device. Then, we validated the model against temperature measurements obtained from a human cadaver skull. Finally, we incorporated skin, brain tissue, and physiological blood flow and quantified their effects on intracochlear temperature during MTH.

### 2.1 Geometry preparation

To develop the model, we constructed human head geometry using anonymized imaging data obtained from the OpenEar Library [31], SICAS Medical Image Repository [32], 3D Slicer library [33], and online opensource DICOM data set of the head [31, 34, 35] (fig 1). The full human head model was obtained from [36], and the internal carotid artery model was downloaded from [37, 38]. The human cadaver skull used in the experiments was scanned using a Revopoint POP 2 structured-light scanner to generate the 3D geometry model required for computational modeling. These images were converted to STL files and imported into COMSOL Multiphysics. Image processing and segmentation was performed using a combination of software including Mimics, 3-Matic, and Blender to ensure compatibility and anatomical accuracy. COMSOL Multiphysics was used to simulate heat transfer and fluid dynamics within the head under MTH, as detailed in the following sections.

**Figure 1:**
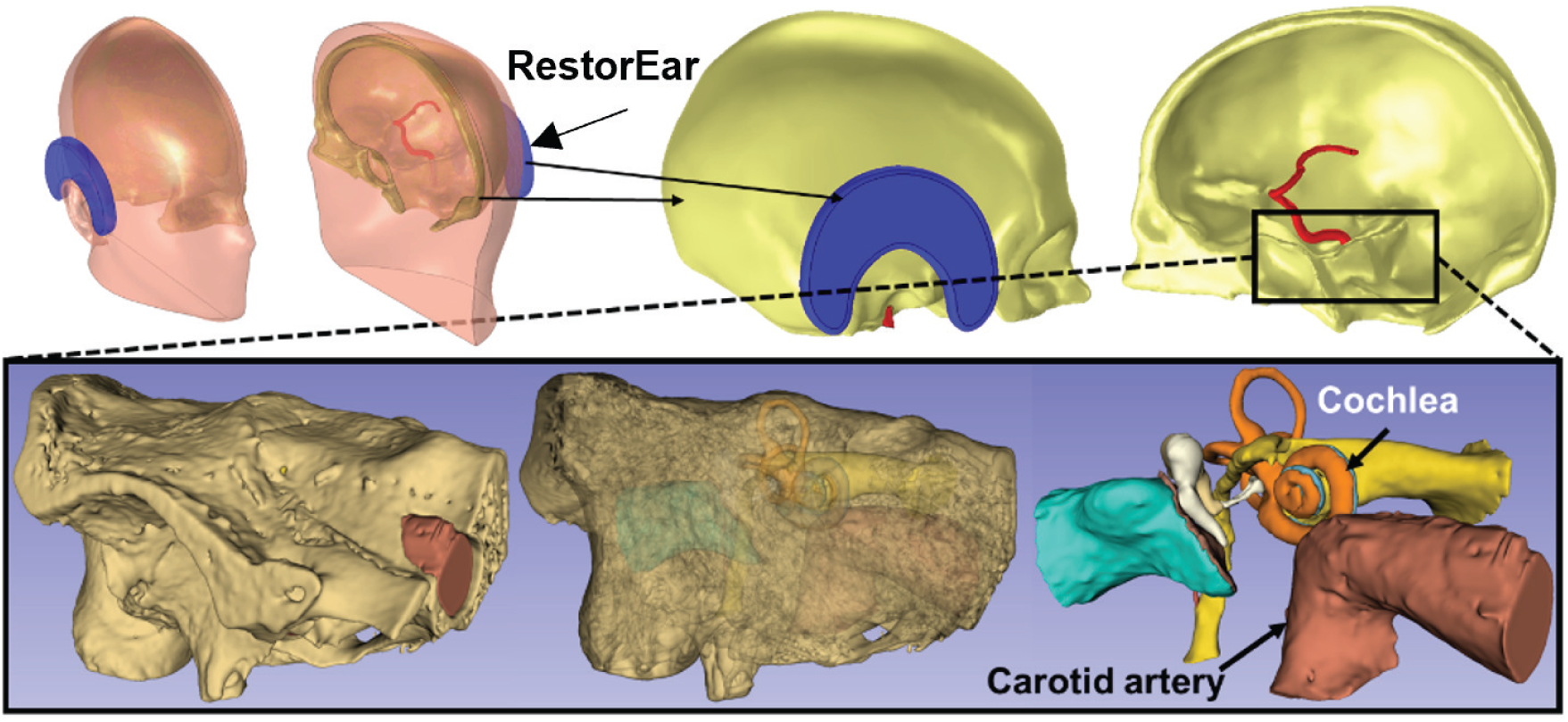
A three-dimensional model of the human temporal bone, obtained through computed tomography and micro-slicing [38] and sourced from Zenodo [34, 31]. Top: One half of a human head with the RestorEar device is illustrated in the image on the left, while the human skull with the RestorEar device and the carotid artery is on the right. Bottom: The temporal bone and a portion of the carotid artery. The middle panel shows a semi-transparent rendering of the cochlea, associated nerves, and carotid artery. The cochlea, middle ear, and carotid artery are displayed in the image on the right. The relative size of the carotid artery compared to the cochlea indicates that vascular flow may play a significant role in thermal therapeutic methods.

### 2.2 Experimental setup

We measured temperature at eight anatomical locations in a human cadaver skull following application of the ReBound Lite MTH device using calibrated thermocouples (Omega, T-type, COCO-005-BW). Temperature drops remained below 10*^◦^*C across all measurements (fig 2). The RestorEar devices consist of gel-based aqua packs designed to induce mild therapeutic hypothermia. These gel packs are stored in a freezer for 24 hours prior to application and undergo passive warming after removal from the freezer. The onset of tissue rewarming was measured to occur approximately 20 minutes after application. Both onset time and temperature drop magnitude were distance-dependent, with later onset and smaller drops recorded in tissue farther from the MTH device.

**Figure 2:**
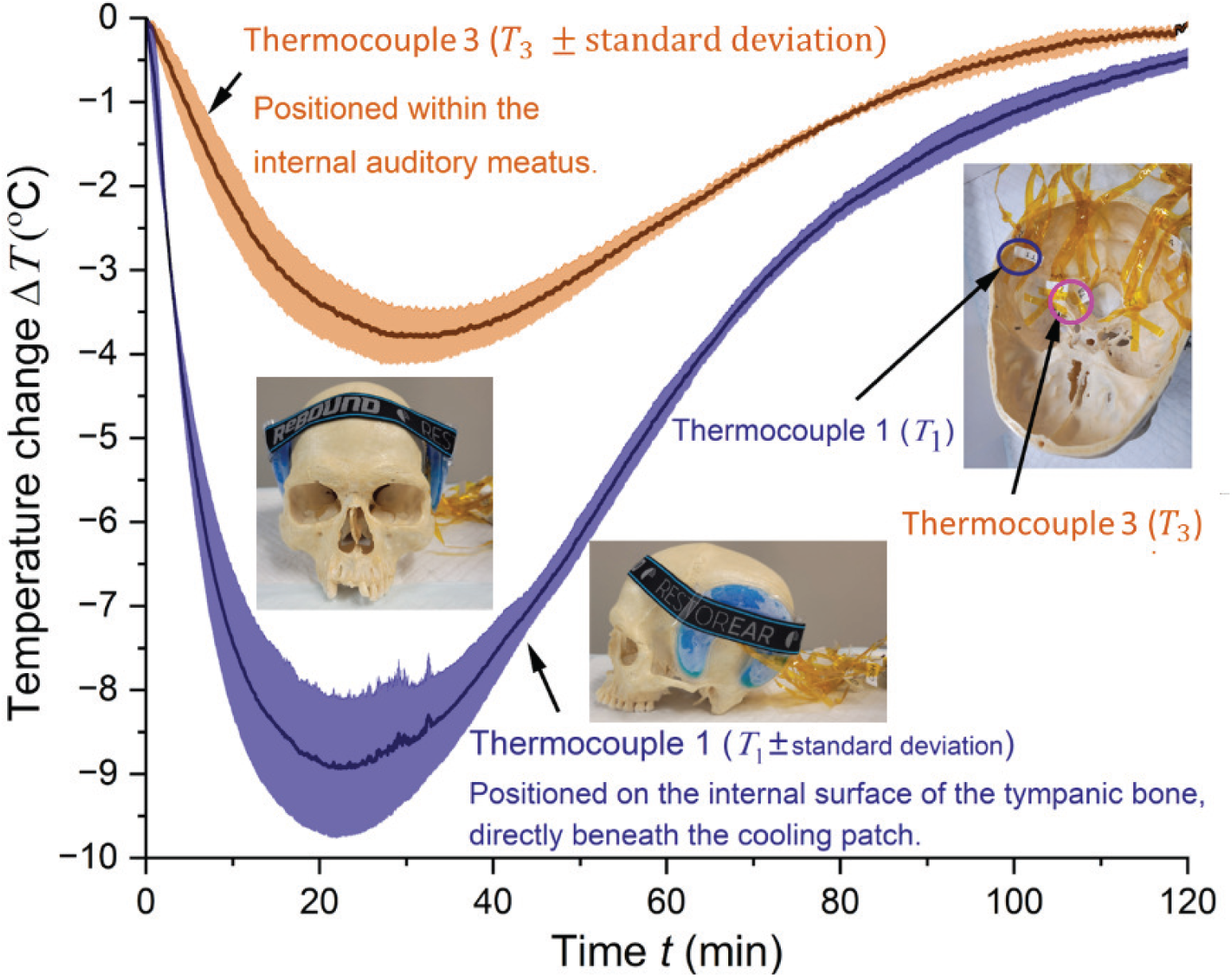
Temperature reduction in a human cadaver skull following application of the ReBound Lite MTH device. Temperature changes were recorded using calibrated thermocouples (Omega, T-type, COCO-005-BW) at eight anatomical locations. Shown in the figure are the maximum and minimum temperature drops, along with their corresponding standard deviations and thermocouple positions. The greatest temperature decrease occurred directly beneath the RestorEar device inside the skull, while the smallest was observed at the internal auditory meatus.

Cadaver skull experiments do not replicate the thermal conditions of the living human ear, where tissue perfusion, metabolism, and physiological heat generation influence temperature distribution. Direct thermocouple measurements in the cochlea of healthy human subjects are not feasible; thus, temperature reduction within the human cochlea was estimated using an in silico model expanded from our previously developed and validated cochlear model [39, 40, 41].

### 2.3 Computational thermal model of cochlear MTH

Heat transfer in human tissues is governed by Pennes’ equation, which includes accumulation, conduction, convection (through perfusion and other biological fluid flows), radiation, metabolic heat generation, and any other heat sources, as stated in Equation 1 [42, 43, 44].

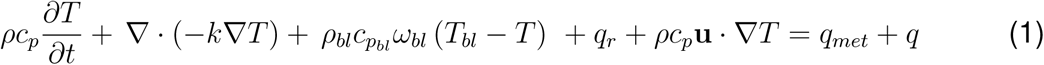

where, *ρ, c_p_, T, t, k, ω, and* **u** are density 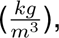 specific heat 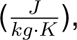 temperature (K), time (s), thermal conductivity 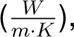 perfusion rate 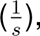 and velocity 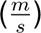 respectively. *q_r_* is the radiative flux 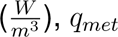 is metabolic heat generation 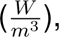 and *q* is power density 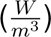 that can be either positive (for heat source) or negative (for heat sink). Radiative flux has a negligible contribution to the overall heat transfer in this study.

The third term on the left-hand side of Eq. 1 ( *ρ_bl_c_p__bl_ ω_bl_*(*T_bl_ − T*)), represents convective heat transfer within the cochlea due to blood flow, while the fifth term (*ρc_p_***u** *· ∇T*), accounts for other forms of convection including the convection due to the movement of cochlear fluids (endolymph and perilymph), and convective heat transfer by blood vessels outside of the cochlea (including internal jugular vein, and internal carotid artery). The perfusion rate 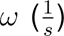 usually is measured empirically and it is an average value. In our model we integrated the perfusion in the convection term (fifth term) to accurately evaluate the heat transfer due to localized perfusion.

The COMSOL Multiphysics material library was used to define the thermophysical properties of the skull, soft tissues, and blood. The cochlear fluids and the MTH cooling device were assumed to have the same thermophysical properties as water (Table 1). The material properties of the MTH device do not affect the simulation results because its transient temperature profile was measured experimentally and used as the model input.

**Table 1:** Thermophysical properties of the materials used in the model.

| Material | $\rho$ ($\frac{kg}{m^3}$) | $c_p$ ($\frac{J}{kg \cdot K}$) | $k$ ($\frac{W}{m \cdot K}$) | $\mu$ (Pa·s) |
| --- | --- | --- | --- | --- |
| Bone | 1908 | 1313 | 0.32 | NA |
| Tissue | 1050 | 3840 | 0.52 | NA |
| Blood | 1080 | 3617 | 0.55 | 0.005 |
| Perilymph, endolymph, cooling gel | 997 | 4182 | 0.60 | NA |

### 2.4 Computational fluid dynamics model of blood flow in cochlea

We hypothesized that the blood flow around the cochlea and the skin surface impacts the effectiveness of thermal therapy by counteracting cooling efforts and maintaining core body temperature. Specifically, the blood flow in vessels near the skin surface decreases in response to cooling (vasoconstriction), minimizing heat loss. As a result, cochlear regions adjacent to these vessels remain warmer than those farther away. To quantify the impact of blood flow on cochlear temperature, we modeled blood flow in the head by solving the momentum balance (Navier-Stokes) equations (Eq. 2) simultaneously with the energy balance equations.

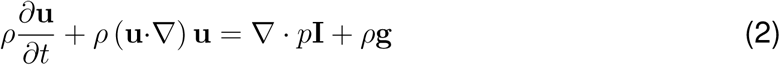

In Eq. 2,the pressure *p* is expressed in pascals (Pa), the gravitational acceleration **g** has units of 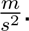 The first and second terms on the right-hand side represent pressure force and body force (gravity), respectively. On the left-hand side, the first term denotes acceleration, and the second term represents advection. This equation was solved for blood flow in internal carotid artery. The resulting velocity was used in Eq. 1 to calculate convective heat transfer. It is well documented that vessels with diameters less than 100*µm* will quickly reach thermal equilibrium with the surrounding tissue and do not affect the temperature of tissues; however, vessels with diameter larger than 500 *µm* will stay at a constant temperature and significantly impact the tissue temperature [45]. The internal carotid artery, the primary blood vessel near the cochlea, has a diameter of several millimeters which is well above the 500 *µm* threshold [46]. Therefore, it is included in this model as a heat source.

### 2.5 Verification and validation

The mesh convergence study was conducted by refining the mesh through three levels of refinement. The temperatures at several critical points were monitored to ensure that the numerical results were independent of the mesh size. Based on the convergence study, the final meshes consisted of 1,070,475 and 2,237,909 domain elements for the skull and full head models, respectively. Additionally, a time-step convergence study was performed using the same procedure, and a time step of 0.1 s was selected for all simulations.

Then we validated the computational model using temperature measurements obtained from a human cadaver skull exposed to an MTH device (fig 2). To replicate the experimental conditions, we simulated the cadaveric skull geometry in COMSOL Multiphysics and calculated the transient temperature distribution during cooling (fig 3). The validated model was subsequently used to investigate the influence of physiological blood perfusion on cochlear temperature during MTH.

**Figure 3:**
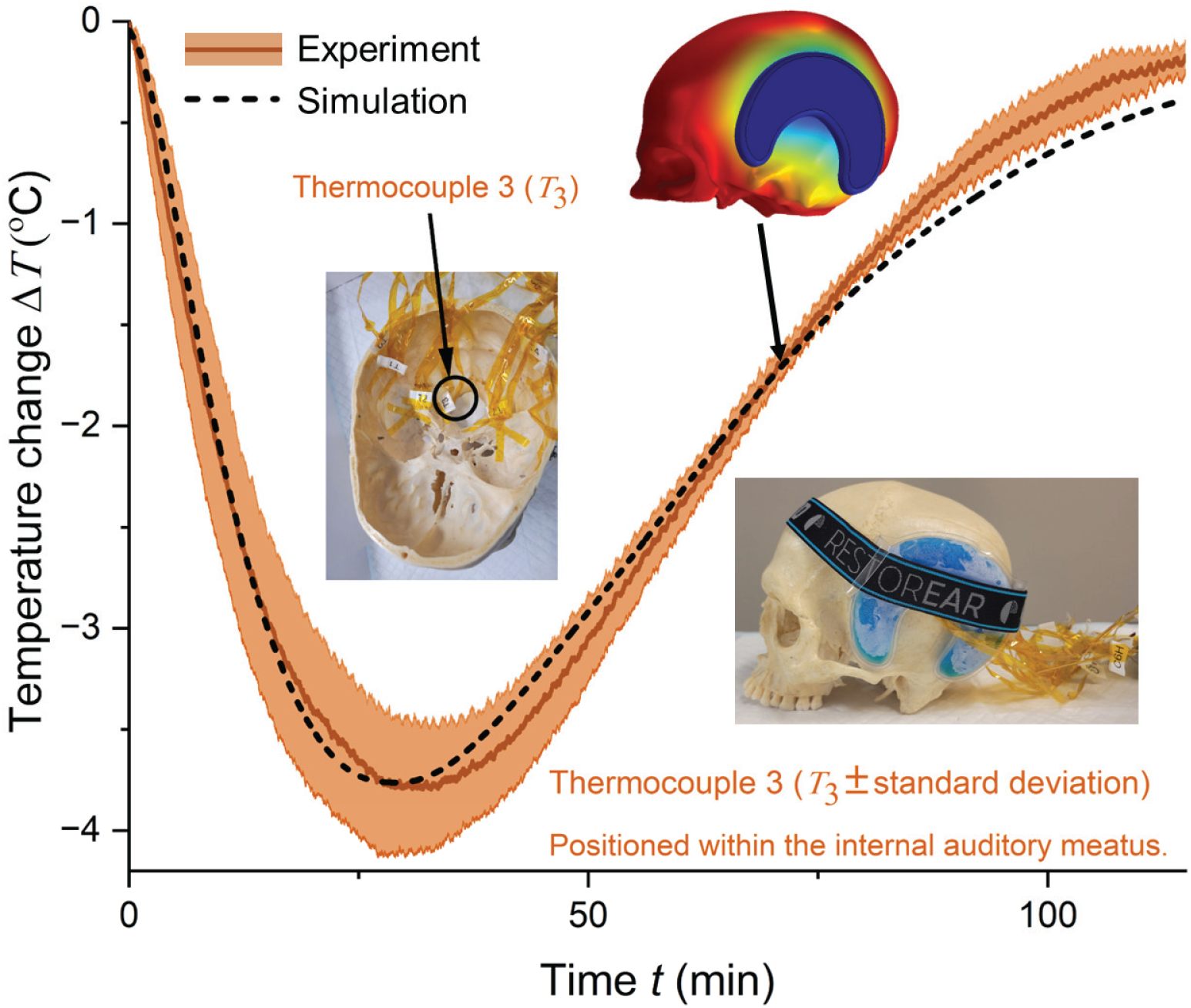
Comparison of the measured and simulated temperatures at the internal auditory meatus following application of the ReBound Lite mild therapeutic hypothermia device to a human cadaver skull. The simulation agreed well with the experimental measurements during the cooling phase, remaining within one standard deviation for approximately the first 75 min. At later times, the simulation slightly underestimated the temperature, likely because changes in the contact between the gel pack and the skull during the melting process were not accounted for in the model.

## 3 Results

We evaluated the impact of an MTH device (ReBoundLite) on human cadaver skull. Recorded temperature reductions peaked at 4*^◦^*C (Δ*T* = *−*4*^◦^*C) at the internal auditory meatus and approximately 10*^◦^*C (Δ*T* = *−*10*^◦^*C) directly beneath the cooling pad on the temporal bone inside the skull (fig 2). This temperature gradient between the location of the RestorEar device on the head and the middle and inner ear was also reported in the recent cadaveric study by Yepes et al. [13]. In that study, the under gel pack thermistor showed a much larger drop than the inner ear (about 24 *^◦^*C, read from their fig 3A [13]). These reported results in reference [13] likely overestimate cooling efficiency, since cadaver studies lack perfusion and vasoconstriction effects, which are crucial determinants of thermal dynamics in vivo. All of these results highlight the importance of considering individual head size, tissue characteristics, and blood perfusion when evaluating MTH protocols and devices.

### 3.1 Validation of in silico model

The temperature within the internal auditory meatus was calculated at the same anatomical location used for the skull temperature measurements, and the experimental results were compared with the corresponding simulation outcomes. The simulation results remained within one standard deviation of the experimental data up to 75 minutes (fig 3). Beyond this time point, deviations were observed, which can be attributed to gel melting, leading to changes in the contact surface area between the gel pack and the skin. This effect was not incorporated into the simulation model.

In the simulation, it was assumed that the gel pack maintained continuous contact with the skull throughout the experiment. Consequently, the simulated temperatures during the last 45 min of rewarming phase were lower than those measured experimentally. To improve model accuracy, dynamic changes in surface contact should be incorporated into future simulations. However, mild therapeutic pre-treatment occurs mainly during the cooling phase; therefore, discrepancies observed during the subsequent rewarming phase may be less critical for this study. This validated model was used to evaluate the effect of blood perfusion on cochlear cooling.

### 3.2 Blood perfusion attenuates cochlear cooling during MTH

We used the validated model to isolate the contribution of vascular heat transport to cochlear cooling. We progressively increased anatomical and physiological realism by first adding the internal carotid artery to the skull geometry (fig 4), and then embedding this enhanced geometry within a full head model that included skin and brain tissues (fig 5). This staged modeling approach provides bounded estimates of cochlear cooling under physiologically realistic in-vivo conditions.

**Figure 4:**
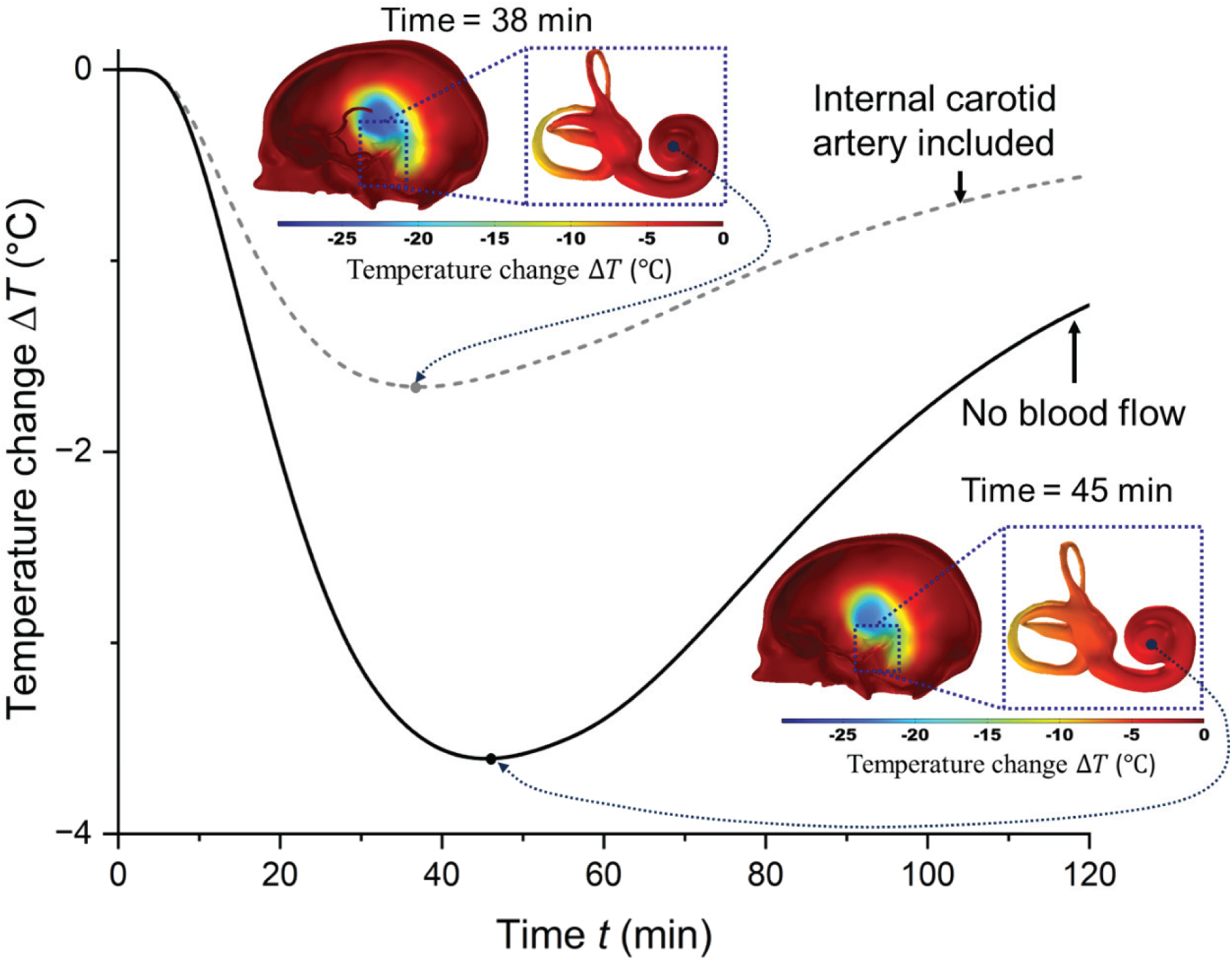
Predicted cochlear temperature changes in the validated skull model with and without internal carotid artery blood flow. Physiological arterial flow (dashed line) limited cochlear cooling to approximately 1.7 *^◦^*C at 38 min after the start of treatment, whereas the absence of arterial flow (solid line) resulted in approximately 3.7 *^◦^*C cooling at 45 min. Insets show cochlear temperature distributions at these minimum-temperature points, highlighting localized warming near the carotid canal due to arterial heat transfer.

**Figure 5:**
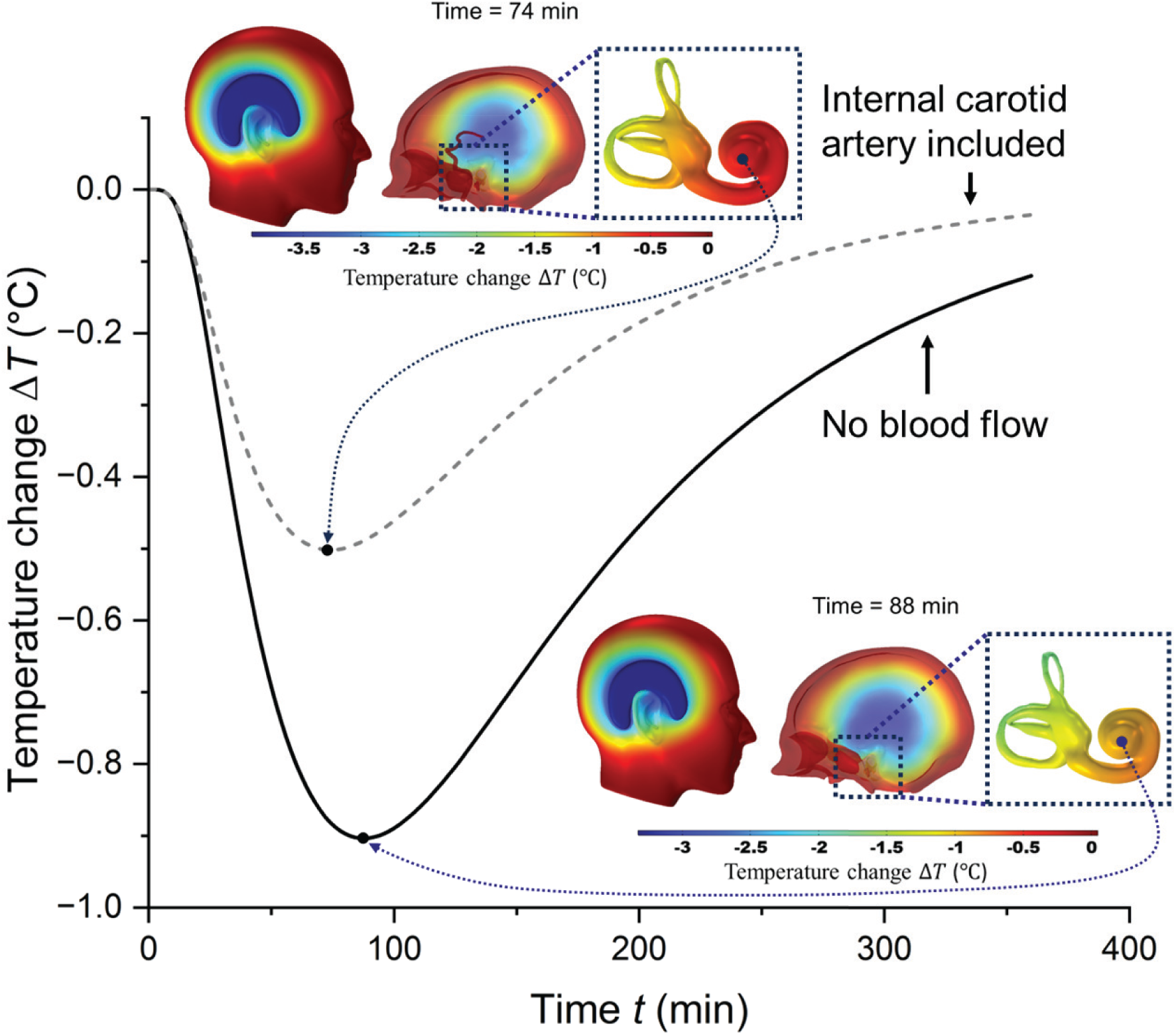
Whole-head simulation of mild therapeutic hypothermia with and without internal carotid artery blood flow. Maximum cochlear cooling is about 0.5 *^◦^*C when arterial flow is included (dotted) and about 0.9 *^◦^*C when blood flow is neglected (solid). Insets show the sagittal head temperature field and the corresponding cochlear distribution, illustrating that soft-tissue conduction and vascular heat transport together limit inner-ear cooling in vivo

### 3.3 Skull-level model with the internal carotid artery

We first introduced physiological blood flow through the internal carotid artery into the validated cadaver-skull model, keeping all other conditions identical to the validation case. The blood inlet temperature and mean velocity were prescribed as physiological values for the internal carotid artery, with values of 37 *^◦^*C and 0.28 m/s, respectively [31]. The experimentally measured RestorEar device temperature was used as the input to the model. The predicted transient cochlear temperature changes in human skull with and without arterial flow, are shown in fig. 4. The transient temperature at the cochlear apex is plotted as a solid black line (without arterial flow) and a dashed gray line (with arterial flow). The temperature distribution profiles shown in the insets correspond to the time points when the minimum cochlear temperature was reached. The minimum temperature occurred at *t*=38 min for the model with the internal carotid artery and at *t*=45 min for the model without the artery. In the absence of the carotid artery, the cochlea cooled by approximately 3.7 *^◦^*C within 45 min. Introducing arterial flow reduced the maximum cochlear cooling to approximately 1.7 *^◦^*C, an attenuation of roughly 2 *^◦^*C (about 54%) (fig 4).

Because the artery continuously delivers heat at core temperature within a few millimeters of the cochlea [43], it behaves as a local thermal source that opposes conductive cooling. This behavior is consistent with classical bioheat theory, in which vessels larger than approximately 500 *µ*m remain close to core temperature and exert a first-order influence on the temperature of the surrounding tissue [37]. Therefore, the carotid artery measurably limits the cooling delivered to the inner ear.

### 3.4 Whole-head model: estimating in-vivo cochlear cooling

Finally, the skull with carotid artery was added within the full head anatomy, and the MTH device was modeled on the skin surface (fig 5). Incorporating the intervening soft tissue and distributed perfusion markedly reduced the cooling that reached the inner ear. The predicted cochlear temperature reduction remained below 1 *^◦^*C, reaching maximum values of 0.5 *^◦^*C with the internal carotid artery included and 0.9 *^◦^*C without arterial flow (fig. 5).

The inclusion of skin thermal resistance and localized blood flow attenuated cochlear cooling by nearly 3 to 4 folds. This highlights that both increased conductive path length and vascular heat transport substantially limit the temperature reduction of the cochlea. Although the magnitude of cooling depends on individual anatomy, device placement, and contact conditions, the qualitative pattern of strong surface cooling that is largely dissipated before reaching the cochlea is expected to be robust. It is important to note that the present model does not include additional insulation from hair or heat exchange with other cranial vessels; thus, the predicted values likely represent an upper bound of achievable cochlear cooling in vivo. Across both models, neglecting the carotid artery systematically led to an overestimation of cochlear cooling, indicating that explicit representation of major vessels and spatially resolved perfusion is necessary to accurately predict MTH device performance in humans. Furthermore, neglecting local blood flow overestimates cochlear cooling duration, which increases from 74 min with perfusion to 88 min when perfusion is neglected.

## 4 Discussions

In this work, we quantified how vascular heat transport constrains localized hypothermia of the human inner ear using a combined experimental and computational framework. Across staged models of increasing anatomical realism, blood perfusion, dominated by the internal carotid artery, kept cochlear temperature close to core body temperature despite pronounced surface cooling. This suggests that non-invasive cooling protocols achieving approximately 6 *^◦^*C cochlear cooling in rodents [30] may not directly translate to humans. In rodents, the inner ear is located closer to the surface and experiences a lower vascular heat load, whereas in the human head, comparable surface cooling results in only a fraction of a degree temperature reduction when the thermal effects of the carotid artery and intervening tissues are accounted for. Our estimates are consistent with the cadaveric measurements of Yepes et al. [13], who reported middle-ear temperature reductions below 1 *^◦^*C after 30 min of cooling in male half-heads and about 3.8 *^◦^*C in female half-heads. The present in silico model is based on a male head. Because cadavers lack perfusion and vasoregulation, those values plausibly represent an upper bound relative to the living head, consistent with the further attenuation predicted here when arterial flow is added.

Methodologically, neglecting localized blood flow systematically overestimates therapeutic cooling. Perfusion averaged forms of the bioheat equation, which distribute a single lumped perfusion term uniformly through the tissue, cannot capture the strong spatially localized heating produced by a large vessel such as the internal carotid artery. Resolving the artery explicitly is essential near highly perfused organs, where a single dominant vessel governs the local thermal balance [37].

This raises a critical question: can MTH still confer meaningful protection when only modest cochlear cooling is achievable in vivo? The protective effects of mild hypothermia may not solely depend on large local temperature reductions. Even small decreases in temperature can reduce metabolic demand, limit excitotoxicity and the generation of reactive oxygen and nitrogen species, suppress apoptosis, and help preserve inner hair cell ribbon synapses [23]. Thus, clinically meaningful protection likely arises from the combination of modest deep cooling and activation of skin temperature sensitive biological processes rather than from deep cooling alone. This shifts the design objective for MTH devices from maximizing temperature reduction to reliably achieving and maintaining a controlled decrease at the target site. Application of non-invasive surface cooling of cochlea is subject to an inherent physiological constraint because the carotid artery buffers cochlear temperature. Therefore, strategies that shorten the thermal conduction pathway or reduce local vascular heat transfer may be more effective than simply increasing cooling power or duration. Furthermore, because the passive gel packs used in this study began to rewarm instantly (fig 2), actively regulated cooling sources that maintain a constant surface temperature may enable more sustained and reproducible therapeutic temperature control.

### 4.1 Study limitations and future perspectives

The current model successfully captures the dominant thermal mechanisms governing cochlear cooling and provides a validated framework for evaluating MTH strategies. Further improvements in model fidelity can be achieved by incorporating additional physiological factors and more comprehensive representations of the human head thermal environment. The present validation was performed using a human cadaver skull, which provided a controlled experimental benchmark for comparing simulated and measured temperatures at the internal auditory meatus. Future validation studies using full-head cadaver models, in-vitro platforms that reproduce the relevant thermophysical properties of the human head, or in-vivo surface tissue temperature measurements could further improve physiological representation. The model currently includes the internal carotid artery but does not include the internal jugular vein, smaller cranial vessels, or dynamic vasoregulation.In addition, tissue properties were assigned using homogeneous literature values, geometry was derived from a single anonymized dataset, device–skin contact was idealized (with contact loss due to gel-pack melting potentially contributing to the deviation observed beyond approximately 75 minutes during validation), and metabolic heat generation and hair insulation were neglected. These assumptions are expected to overestimate cochlear cooling.

Future work will incorporate more detailed vascular networks and thermoregulatory responses, include subject-specific anatomy derived from imaging, and couple thermal predictions with biochemical models of hair-cell protection to establish relationships between temperature exposure and functional outcomes. Incorporating in vivo human temperature and blood perfusion data would further complete the validation and verification process, providing a more comprehensive assessment of the model. More broadly, the validated framework provides a generalizable approach for studying localized thermal therapies near highly perfused tissues, where vascular heat transport critically influences achievable temperature reduction.

In summary, vascular heat transport imposes a physiological ceiling on achievable cochlear cooling in humans. The internal carotid artery limits the temperature reduction reaching the cochlea to well under 1 *^◦^*C despite pronounced surface cooling. This limit suggests that the therapeutic benefit of MTH for hearing preservation, now being tested in first-in-human trials (ClinicalTrials.gov NCT06375278 and NCT06729632), likely derives from temperature-sensitive protective pathways triggered by modest cooling rather than from deep hypothermia, and it underscores the value of the validated framework presented here for guiding future device design and cross-species translation.

## Acknowledgement

We thank Dr. Rehana Lovely (UNT Health-Center for anatomical sciences, College of Biomedical and Translational Sciences, Physiology and Anatomy Department) for providing the human skull used in the experiments. We acknowledge the University of North Texas for faculty start-up funding, two seed grants that supported this research, and NIH 1R15DC023030-01A1.

## 5 Statements and Declarations

### Ethics Approval

This study used a human cadaver skull. U.S. federal human-subjects regulations (45 CFR 46) define human subjects research as involving living individuals; because decedents do not meet this definition, Institutional Review Board approval was not required for this work.

### Data Availability

The datasets generated during this study are publicly available in the GitHub repository at https://github.com/TFAM-Lab/VascularHeat-MTH.git.

### Author Contributions

*B. McCorkendale*: Data collection, manuscript preparation, and approval of the final manuscript.

*R. Rodriguez, R. Fink, and M. Moore*: Data collection and approval of the final manuscript.

*S. A. Romero*: Study conception, data analysis, manuscript preparation, and approval of the final manuscript.

*F. Esmailie*: Study conception, study design, data collection, data analysis, manuscript preparation, and approval of the final manuscript.

